# Tailoring a glycerol-free cryopreservation protocol with anti-freeze (glycol)proteins for commercial and native breeds of chicken

**DOI:** 10.1101/2025.07.18.664799

**Authors:** Berenice Bernal, Éva Váradi, Árpád Drobnyák, Tim Hogervorst, Krisztina Liptói, I. K. Voets, Agnes de Wit, Julián Santiago-Moreno, Rachel Hawken, Carolien de Kovel, Sipke-Joost Hiemstra, Henri Woelders, Barbara Végi

**Author notes:** Corresponding author: Agnes de Wit, Centre of Genetic resources, the Netherlands. Wageningen University and Research, 6700 AH, Wageningen, the Netherlands. aEmail address.

## Abstract

This study evaluated the effects of antifreeze proteins AFPI, AFPIII, and AFGP during fresh processing, and freezing-thawing of chicken semen from a commercial broiler breed (BB) and the local breed Yellow Hungarian (YH), using 0.6 mol/L DMA as cryoprotectant. AFPI had small but significant (p<0.05) positive effects on post-thaw sperm viability and motility in BB. For YH semen, three freezing protocols with different dilution rates (4×, 2.3×, 2.1×) were compared. Protocol 1 (dilution rate 4×) gave the best post-thaw viability (P<0.001) and was used further. Fertility rates (FR) of pre-freezing steps were tested by inseminating WL hens (4 AI/hen in 2 weeks, 100 × 10L sp/dose). FRs for A) raw semen, B) chilled semen, C) B + DMA, D) B + AFPI, and E) B + DMA + AFPI were 48.2, 3.7, 31.4, 21.4, and 16.1%, respectively. B was lower than A (p<0.001), while C–E were higher than B (p<0.005). AFPI during freezing gave no advantage compared with DMA, except for improved post-thaw DNA integrity. Inseminations with frozen-thawed semen (8 AI/hen in 3 weeks, 100 × 10L sp/dose) gave low FR with YH semen, regardless of hen type. FR with YH semen was 0% without AFPI and 1.5% with AFPI. The oviduct embryo mortality (18.0%±1.4) observed across all the different hen groups suggested insufficient number of spermatozoa inseminated. In conclusion, AFPI improved some post-thaw traits, but fertility outcomes remain inconclusive. Preventing pre-freezing fertility loss and increasing sperm dose concentration are required.

## 1. Introduction

Sperm cryopreservation plays an important role in ex-situ conservation of genetic diversity. However, loss of cell functional intactness by ice formation is a major obstacle in many livestock species. Currently, the dose of cryopreserved chicken semen required for artificial insemination (AI) is 50 times higher than that of fresh semen [1] and even with such high sperm dosage, the fertilization rates with frozen-thaw semen remain variable and poor for many breeds/lines of chickens [1–4]. Moreover, the demonstrated dependence between cryotolerance and chicken breed [5] seems to indicate the need to adapt cryopreservation protocols to the breed of interest. Thus, research continues evolving regarding chicken sperm cryopreservation.

The freezing of a tissue or a suspension of cells leads to cryoinjury and cell death. Much of this is related to the formation of ice and the resulting osmotic stress [6]. Intracellular ice formation is considered absolutely lethal and can be prevented by allowing extracellular ice formation at optimal cooling rates [3,6,7]. However, the large extracellular ice masses and ice crystals can physically damage cells. This damage can vary depending on the form and size of ice crystals which is affected, amongst other factors, by recrystallization, i.e., the change in the size, shape or orientation of individual crystals after the completion of solidification [8].

Antifreeze proteins (e.g. AFPI, AFPIII, AFGP) are found in freeze-avoiding and freeze-tolerant organisms (animals, plants, fungi, or bacteria). These proteins are capable to bind to the surface of nascent ice crystals [9,10] and control the size, shape and aggregation of smaller ice crystals, thus affecting the mechanical stress and cell damage caused by ice [11,12]. At micromolar concentrations, they inhibit ice recrystallization, [13], and it is proposed they interact with the plasma membrane [14,15]. Another unique property is its ability to reduce the freezing point of solutions (thermal hysteresis) in a non-colligative manner, allowing this effect to be achieved using significantly lower amounts compared to traditional cryoprotectants [15].

AF(G)Ps can vary according to their composition, structure, size, mode of action, morphology of ice crystals produced and ice recrystallization inhibition (IRI) activity. The present study focused on studying AFPI from Winter Flounder (Pseudopleuronectes americanus), AFPIII from Ocean Pout (Zoarces americanus) and a synthetic AFGP containing the native disaccharide (sequence: AATdAATdAATdAATdAA).

AFPIs, are 3-4.5 kDa alanine-rich α-helice proteins capable to bind to the pyramidal faces of ice, while AFPIIIs are small globular protein of 6.5-14 kDa that bind to both, the prism plane and the pyramidal plane of ice [12,16]. AFGPs are 2.7–32 kDa proteins formed of 4 to 50 tripeptide repeats of Ala–Ala–Thr with the disaccharide galactose-N acetylgalactosamine attached to each Thr [12,16] and they bind to the prism plane of ice crystals [17].

In contrast to AFPs that can be obtained by recombinant protein expression techniques, the glycosylation pattern on AFGPs cannot be obtained via protein expression and therefore pure samples of defined length are currently only obtained via chemical synthesis [15].

AF(G)Ps have been used in sperm cryopreservation of different animals such as: rabbit [18], buffalo [19], ram [20], miniature pig [21], sterlet [22], among others [23] observing positive effects in some cases but non or detrimental effects in others. The kind of effect seems to depend on various factors such as kind and concentration of AF(G)P, the species, and also the cryopreservation protocol used [23].

Regarding chicken, a work done with AFPIII to freeze sperm from Ross 308 breeder-breed roosters [24] reported improved post-thawed sperm quality and fertility by supplementing 1 mg/mL of the protein in the freezing media. Another study conducted with White Leghorn chicken sperm, supplemented with winter wheat AFGP [25], revealed an increase in fertility with frozen-thawed semen from 31- and 65-week-old roosters, using 0.1 µg/mL and 1 µg/mL of the protein, respectively. These protocols employed glycerol as cryoprotectant, which is known to have a contraceptive effect in hen, thereby requiring its removal prior AI. In this context, the present study investigated the effects of AFPI, AFP III, and AFGP on the freezability of chicken sperm using DMA as cryoprotectant.

A previously validated cryopreservation protocol, that demonstrated improved post-thaw semen quality in broiler-breeder roosters using DMA as cryoprotectant [26], was taken as a base. Furthermore, the protocol was tested on a native chicken breed (Yellow Hungarian), given that indigenous breeds often exhibit increased sensitivity to cryoinjury and thus serve as suitable models for the optimization of semen cryopreservation strategies.

## 2. Materials and methods

### 2.1 Experimental Design

The experimental design of the present work consisted of two parts (Fig. 1). Part 1 was carried out at the Centre for Genetic Resources, the Netherlands, in Wageningen University and Research (WUR) and the Eindhoven University of Technology (TUE), in Netherlands. This part involved the *in vitro* evaluation of AFP I, AFPIII and AFGP (synthetized by the group of I. Voets, TUE) to select the best type of protein, and its respective concentration, to increase the cryotolerance of chicken sperm. The range of concentrations evaluated was established based on IRI activity tests (described below, Fig. 2A, 2B). The freezing experiments with the different AF(G)Ps were carried out with sperm of commercial broiler-breed roosters (BB) of the company Cobb (the Netherlands). Part 2 involved three experiments to evaluate the effect of the selected AFP on semen from the native breed Yellow Hungarian (YH), at the National Centre for Biodiversity and Genetic Conservation (NBGK), Hungary. The first freezing experiment was performed without AFP to compare 3 different freezing protocols and select the best one for this breed. Once selected the best freezing protocol, pre-freezing and freezing experiments (experiments 2 and 3) were performed to evaluate the effect of supplementing AFPI in the poultry extender. Both experiments included *in vitro* tests and fertility rate evaluations by artificial insemination (AI).

**Fig. 1.**
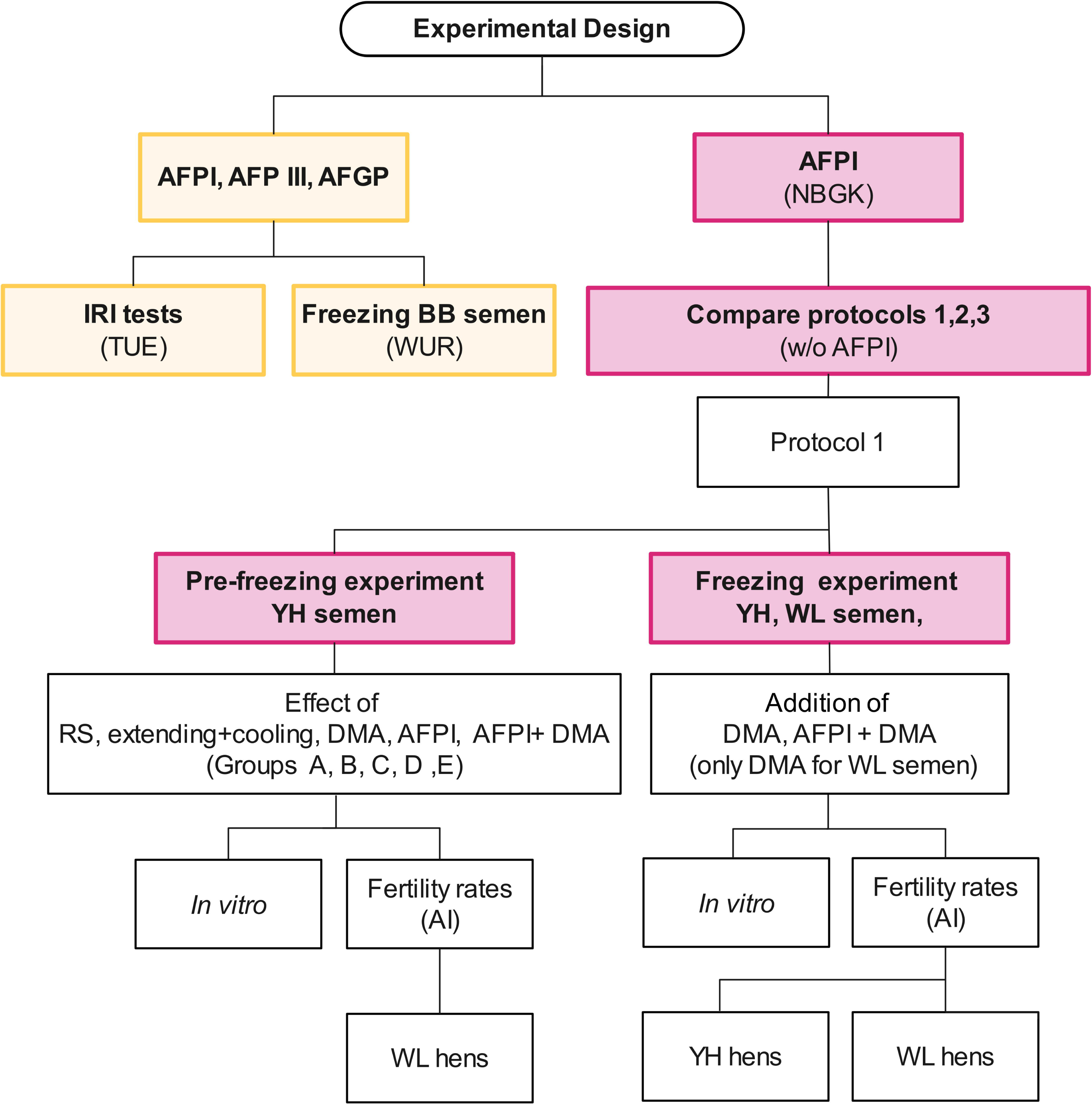
Experimental Design. AFPI, antifreeze protein I; AFPIII, antifreeze protein III; AFGP, antifreeze glycoprotein; IRI, ice recrystallization inhibition; BB, broiler-breed; TUE, Eindhoven University of Technology; WUR, Wageningen University and Research; NBGK, National Centre for Biodiversity and Gene Conservation (Hungary); w/o, without; YH, Yellow Hungarian; WL, White Leghorn; DMA, dimethylacetamide; AI, artificial insemination.

**Fig. 2A.**
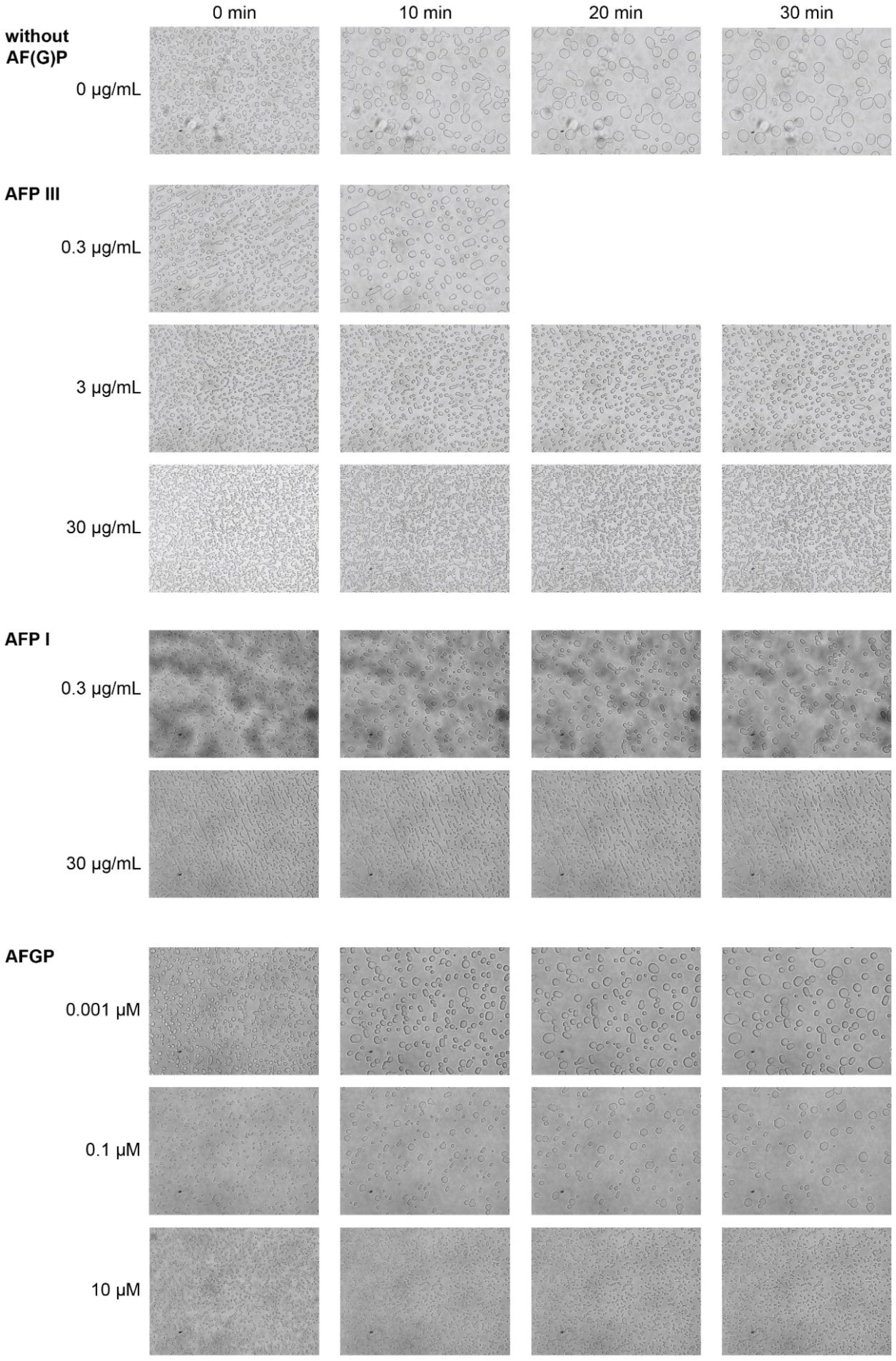
Microphotographs of ice recrystalization activity assay for extender + 1.8 mol/l of DMA (top, 0 ug/mL), and dilutions of AFP III (0.3, 3 and 30 ug/mL), AFP I (0.3 and 30 ug/mL) and AFGP (0.001, 0.1, 10 µM).

**Fig. 2B.**
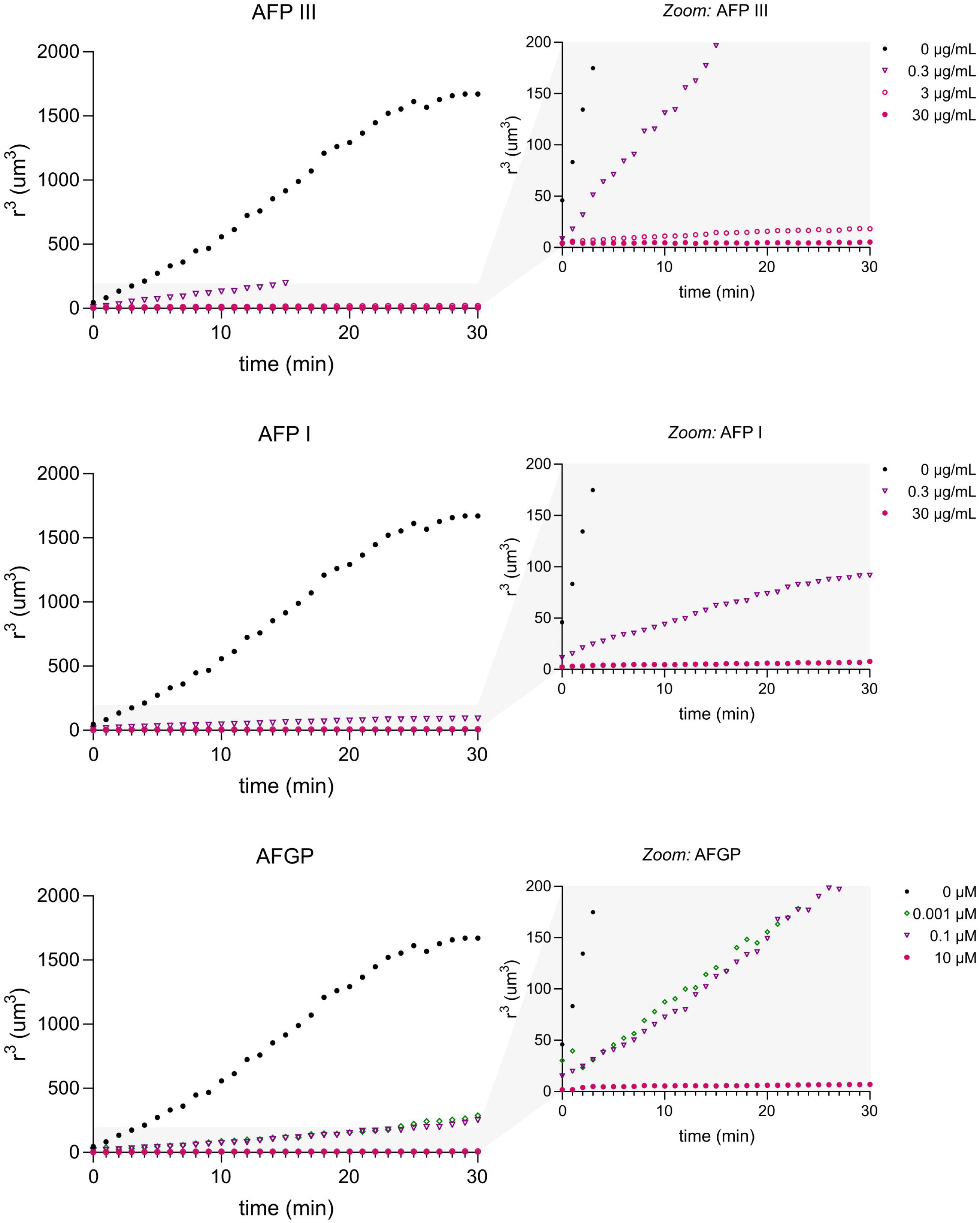
Average cubic radio of ice crystals in extender + DMA 1.8 mol/l over time, in the pressence of AFP III, AFP I and AFGP at different concentrations.

### 2.2 ASG-PE extender

Animal Sciences Group poultry extender (ASG-PE) was used in the present work. This extender contained (gram per 100 ml, mmol/l between brackets): 1.21 g (64.7) sodium-L-glutamate.H2O, 0.102 g (3.14) tri-potassium-citrate.H2O, 0.064 g (2.97) magnesium acetate.4H2O, 0.53 g (26.5) D-(+)-glucose.H_2_O, 2.43 g (114) BES (N,N-Bis (2-hydroxyethyl)-2-aminoethanesulfonic acid), 0.185 g (46.2) NaOH, and Milli-Q water to a final volume of 100 ml, resulting in pH 7.1 and osmolality of approximately 325 mOsm/kg of water [26].

### 2.3 Ice recrystallization activity of AF(G)Ps

For each of AF(G)Ps the IRI activity was screened in ASG-PE containing DMA 1.8M at TUE. These assays where performed to assess whether the components of the poultry extender would affect the IRI activity of the AF(G)Ps and to identify at what concentration AF(G)Ps would still induce IRI activity. Dilutions of AF(G)Ps in extender containing DMA were sandwiched between two precleaned 22x22 mm cover slides and rapidly cooled to −40 °C (20 °C/min), reheated to −10 °C (10 °C/min), to −8°C (1 °C/min), and held at −8 °C for 30 minutes in a Linkam LTS420 stage attached to a Nikon ECLIPSE Ci-Pol Optical Microscope with a Lumenera INFINITY 3 CCD camera. Microphotographs were taken every minute (Figure 2A), from which ice crystal areas were extracted using ImageJ and converted into average cubic radii that was plotted as a function of time assuming a circular crystal shape (Figure 2B).

### 2.4 Semen collection

Semen was collected by the abdominal massage technique [27] and was diluted 1:1 (v:v) directly (in the barn) after collection with ASG-PE at room temperature and transported to the laboratory.

### 2.5 Animals and semen processing

#### 2.5.1 CGN (WUR)

Semen was kindly provided by Cobb from a local research facility in the Netherlands. The semen was obtained from broiler-breeder cocks at the age of 33–39 weeks. The samples were transported to the laboratory of CGN in approximately 30-45 min in a cooling box at 5 °C. Once in the laboratory, the semen samples were pooled (5 roosters per pool) and further processed in a refrigerated work bench set at 5 °C. Motility and viability of each pool were evaluated before addition of the cryoprotectant.

For freezing, mother solutions of AFPI, AFPIII and AFGP were prepared at concentrations of 180 µg/mL, 60 µg/mL, 60 µM, respectively with ASG-PE. From these solutions the corresponding dilutions were done with ASG-PE to obtain solution at concentrations of 180,18, 6, 1.8 and 0 µg/mL of AFPI, 60, 6, 0.6, 0.06 and 0 µg/mL of AFPIII and 60, 6, 0.6, 0.06 and 0 µM of AFGP. Then, 0.333 mL of these solutions, pre-cooled at 5 °C, were added to 1 mL of pooled semen followed by addition of 0.666 ml of ASG-PE containing DMA 1.8 M and gentle mixing (final semen dilution ratio [**DR**] 4x [1:3 v:v]). The final concentration of DMA was 0.6M and 30, 3, 1, 0.3 and 0 µg/mL of AFPI; 10,1,0.1, 0.01 and 0 µg/mL of AFPIII; and 10, 1, 0.1, 0.01 and 0 µM of AFGP. The pools were held with DMA and the different concentrations of AF(G)P at 5°C for 1h, and then packed in 0.5 French straws. The straws were placed in an automatic freezer (Ice Cube, 14S, Minitube, Tiefenbach, Germany) and frozen in three steps: from 5 °C to –20 °C at a rate of –50 °C/min, followed by a 20-second hold at –20 °C, and then continued from –20 °C to –140 °C at the same rate. This protocol follows the recommendation by Woelders et al. (2022) [26] to achieve a cooling rate (CR) of ≥ –50 °C/min. The frozen straws were then plunged into and maintained in liquid nitrogen (at −196 °C) until thawing. For thawing, the straws were warmed for 30 seconds in an agitated water bath at 5 °C. Sperm quality was evaluated.

#### 2.5.2 NBGK

Fifteen Yellow Hungarian (YH) roosters and fifteen White Leghorn (WL) roosters of 1-year-old were housed under 14L:10D photoperiod and natural temperature conditions in individual deep floor cages at the National Centre for Biodiversity and Gene Conservation, Gödöllő, Hungary. All birds had a constant diet throughout the experimental period. Specifically, the birds were fed with commercial feed containing 16,3 % crude protein, 10,95 AME/MJ, 3,3% crude fat, 4,2% crude fibre, 13,7 crude ashes

##### 2.5.2.1 Freezing protocols without AFPI

A first experiment was performed in absence of AFP to find the best protocol for YH sperm (YH-sp). Three freezing protocols were evaluated.

*Protocol 1.* One mL of semen was diluted with one mL of ASG-PE (DR 1:1) at room temperature in the barn. The samples were transported to the laboratory in approximately 10 min and placed in a cooling hood at 5 °C. After 10 min, the samples were diluted for second time with 0.666 mL of ASG-PE and 1.332 mL of ASG-PE containing 1.8 M of DMA (final DR 4x [1:3 v:v] and final concentration of DMA 0.6 M). The samples were packed in 0.5 mL straws and placed on a polystyrene rack of 2 cm height. The rack was then placed on the surface of liquid nitrogen inside a polystyrene box with internal dimensions of 44.3 cm × 26.8 cm × 20.8 cm (L × W× H), which had been filled to a height of 6 cm (equivalent to 5.53LL) with liquid nitrogen. After 10 min, the straws were plunged directly in the liquid nitrogen and stored. The cooling rate achieved with this protocol was ≈ –100L°C/min) which follows Woelders et al. (2022) recommendations

*Protocol 2* was similar to *Protocol 1* with the difference of diluting 1 mL of semen with 0.5 mL of ASG-PE (DR 1:0.5, v:v) in the barn. Once in the lab and after 10 min of storage at 5 °C, a volume of 0.75 mL of ASG-PE containing 1.8 M of DMA was added (final DR 2.3x [1:1.3 v:v], final concentration of DMA 0.6M). The diluted sperm samples were kept for 1h at 5° C and frozen as in *Protocol 1*.

*Protocol 3.* Protocols 1 and 2 were compared with the standard method used for YH-sp (*Protocol 3*). Semen was diluted with ASG-PE at room temperature in the barn (1:1, v:v), taken to the lab in 10 min and placed in a cooling hood at 5 °C. After 10 min, DMA (6% of the semen volume, approximately 0.65 mol/l) was added to the extended semen (final DR 2.1x [1:1.1, v:v]) and the samples were maintained at 5 °C for 30 minutes. The samples where then packed in 0.25 mL straws. Freezing was performed in two steps: first, by placing the straws 5 cm above the surface of liquid nitrogen for 15 minutes; then, at 1 cm above the surface for additional 15 minutes.

##### 2.5.2.2 Pre-freezing experiment with/without AFPI

The effect of each pre-freezing step on sperm quality and fertility were evaluated. These steps were: semen extension + cooling (B), addition of DMA (C) or AFPI (D), and AFPI + DMA (E). Each step was evaluated *in vitro* (motility, sperm abnormalities, viability, intact sperm, acrosome integrity and DNA integrity) and by artificial insemination (AI). Raw pooled semen (10 randomly selected YH roosters / pool) was divided to evaluate steps B-E (RS for B-E). Another, unprocessed pool of semen (5 randomly selected YH roosters / pool) was used to evaluate the quality and fertility of raw YH-sp (step A, RS for AI). Four inseminations were performed (twice a week/ two weeks) using the semen from each pre-freezing step (100 million YH-sp/hen). Five groups of White Leghorn (WL) hens (10 hens/group) were inseminated with: A (raw semen, RS for AI), B (raw semen diluted 4x with ASG-PE and chilled for 1h at 5°C), C (B+DMA), D (B+AFPI) and E (B+DMA+AFPI). Collection of eggs, started from day 2 after the first insemination until day 3 after the last insemination. Fertility (% fertile/incubated eggs) of all the inseminated groups was determined by candling after 7 days of incubation.

##### 2.5.2.2 Freezing experiment with/without AFPI

Cryopreservation of semen with AFPI was done as Protocol 1 with the exception of adding, in the second dilution, 0.666 mL of ASG-PE containing 6 µg/mL of AFPI, (instead of ASG-PE alone). Cryopreservation of YH-sp and WL-sp without AFPI were used as reference. The effect of AFPI on frozen-thawed sperm was evaluated *in vitro* (motility, sperm abnormalities, viability, intact sperm, acrosome integrity and DNA integrity) and by AI. Fifty hens (20 YH and 30 WL) were divided into 5 groups (n=10/group) and inseminated with different kind of semen: (1) Raw YH-sp - WL hens; (2) F/T YH-sp - WL hens; (3) F/T YH-sp - YH hens; (4) F/T YH-sp + AFPI - WL hens, (5) F/T WL semen - YH hens and (6) F/T WL semen - WL hens. Semen for Groups 2, and 4 were frozen in a split-sample approach. Hens were inseminated with ≈100 million spermatozoa as follows: 3 AI/week for 2 weeks, and 2 AIs on the third week (8 inseminations in total). Group 1 was used to know fertility of raw semen of the YH males. Comparisons (1) vs (2) assessed the effect of freezing (without AFPI) on YH-sp; (2) vs (4) assessed the effect of supplementing AFPI to the freezing medium; (2) vs. (3) and (5) vs (6) aimed to detect influence of the female breed when using F/T semen and (3) vs (5) and (2) vs (6) aimed to detect influence of the males breed when using F/T semen. Fertility was determined as described in the pre-freezing experiment.

### 2.6 Assessment methods

*Sperm concentration and motility.* Sperm concentration was determined using an Accucell (IMV, L’Aigle, France) at WUR and NBGK. Motility was assayed using a computer-aided sperm analyzer system consisting on a camera (BASLER avA1000-100gc, Germany[WUR] / BASLER ace U acA1300-200uc, Germany [NBGK]) coupled to a phase contrast microscope (Zeis Axioscope A.1, Germany [WUR] / Nikon Eclipse E200, Japan [NBGK]), and using the 12500/0000 Androvision^®^ software (Minitübe, Germany) [WUR] / Sperm Class Analyzer (SCA; Barcelona, Spain) v.6.0. Software (Microptic S.L.) [NBGK]. For motility analysis, sperm samples were diluted to a concentration of approximately 40 million sperm/mL and loaded onto glass slide at room temperature (WUR) or 37 °C (NBGK). The proportion of motile spermatozoa and the proportion percentage showing progressive motility were recorded. Sperm kinematic variables – velocity (VCL), straight-line velocity (VSL), average path velocity (VAP) – were also recorded. It has been shown that VSL and VAP are the two variables most related to the ability of the sperm to reach the sperm storage tubules (SSTs) at the utero-vaginal junction and subsequent fertility [28]. A minimum of 5 fields and 200 sperm were evaluated at a magnification of 100x for each sample.

*Viability* of the samples, at WUR facilities, were determined by DAPI (4′,6-Diamidino-2-Phenylindole, Dihydro-chloride) diluted in ASG-PE. A sperm aliquot of 20 μl of semen was mixed with 20 μl of 10 μmol/L DAPI, held for 5 min at room temperature in the dark, and then mixed with 10 μl of glutaraldehyde (0.5 vol%) to immobilize the sperm. Two-hundred cells per sample were evaluated by epifluorescence microscopy (Zeis Axioscope A.1, Germany). The percentage of sperm cells that excluded DAPI, i.e. cells with membrane integrity was determined [26]. At NBGK, viability was determined using propidium iodide and SYBR-14 [29]. Two hundred cells were examined under an epifluorescence microscope (Zeiss Axioskop 2 plus, Austria) at 40× objective (wavelength: 450–490 nm).

*Morphological abnormalities* (Abn) were assessed by eosin-aniline staining. The eosin-aniline staining was prepared by dissolving 0.2 g of eosin and 0.8 g of aniline in 10 mL of Lake extender. Twenty µl of the prepared dye were pipetted into an Eppendorf tube and mixed with 10 µl of semen. Ten µl of the mixture were placed onto a microscope slide, spread evenly and fix the sample using a hairdryer, holding it approximately 20 cm away to avoid overheating. The smears were examined under an oil immersion objective at 1200× magnification (Zeiss Axioskope microscope, Germany). A total of 200 sperm cells were evaluated per slide [30]. The proportion of intact live sperm (ILS) was determined.

*Acrosome integrity* (AcI). Twenty microliters of semen were transferred to vials containing 20 µl of formaldehyde 4% in PBS for fixation. The samples were left in formaldehyde for 30 min at room temperature (for non-chilled samples) and 5 °C (for chilled and post-thaw samples). Ten µL drops of the fixed samples (chilled/FT and warmed) were spread onto glass slides (smears) and allowed to dry. For evaluating the acrosome integrity (AcI), the smears of fixed sperm were immersed in a solution of methyl blue 2.5% in PBS for 5 min, then washed with distillated water and let dry. The methyl blue preparation was adapted from Santiago Moreno et al. (2009) [31]. Briefly, methyl blue (Sigma-Alrich) was dissolved in PBS to a concentration of 2.5% and the pH adjusted to 3.5 with acetic acid 2%. The slides where mounted and analysed under a phase contrast microscope (Zeiss Axioskop 2 plus, camera Micro Publisher 6, Canada) at 1000× magnification. Sperm with abnormal morphology of the acrosome (hooks, swollen, thinned or absence) were considered lacking in acrosome integrity [31]; 200 cells were examined.

*DNA integrity* was assessed as described by Bernal et al. (2021) [32] and the samples analysed under a fluorescent microscope (Zeiss Axioskop 2 plus, camera Micro Publisher 6, Canada). Percentages of positive TUNEL spermatozoa per sample were recorded by counting a minimum of 200 spermatozoa per microscopy preparation.

### 2.7 Identification of very early (oviductal) embryo death

After the eggs were cracked, yolks were carefully separated from the albumen and placed into a 0.9% NaCl solution. Germinal discs that appeared infertile were gently removed from the vitelline membrane, immersed in 0.9% NaCl, and stained on microscope slides with propidium iodide (PI). The working solution contained 5 µg PI per ml of 0.9% NaCl, from which 5 µl was applied to each slide. The presence of cell nuclei, serving as an indicator of fertility (Fig. 3), was assessed using a fluorescence microscope (Leitz-Diaplan) at 500× magnification [33].

**Fig. 3.**
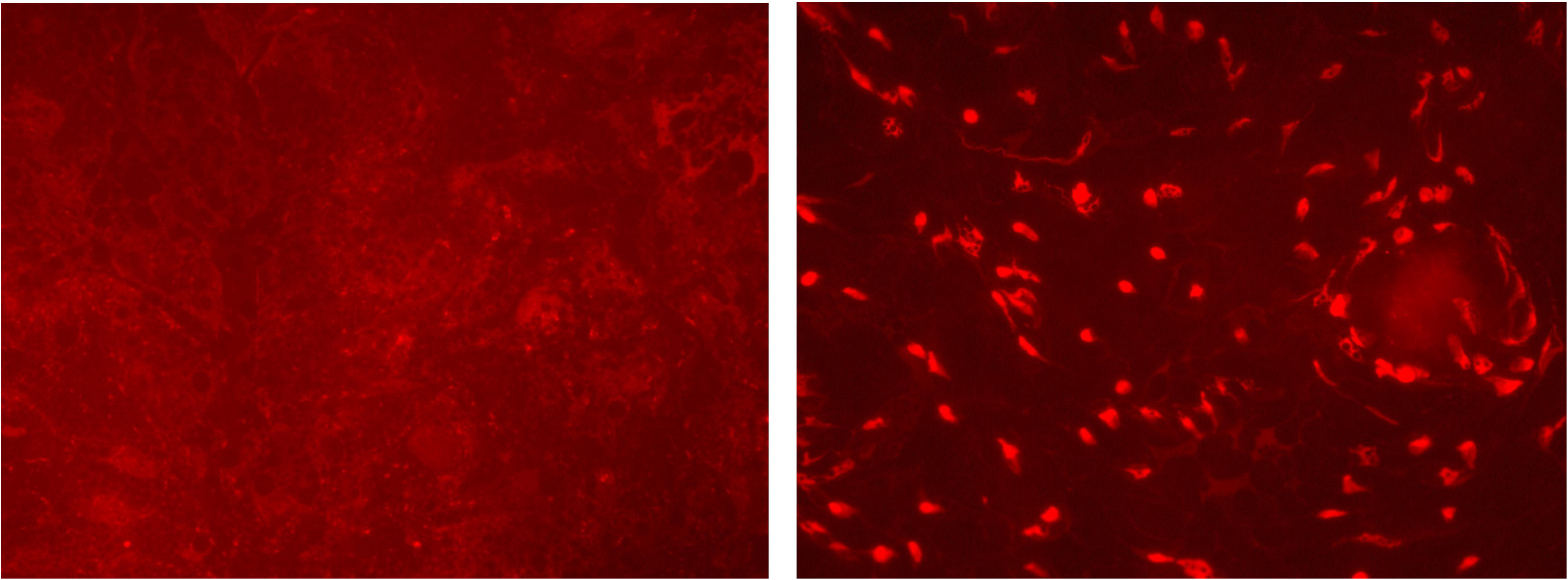
Identification of very early (oviductal) embryo death. Germinal discs were gently removed from the vitelline membrane, immersed in 0.96 % NaCl and stained on microscope slides with propidium iodide. The presence of cell nuclei is an indicator of fertility. Unfertile germinal disc (left) and fertile germinal disc (right). (Photographs taken by Dr. Krisztina Liptói, NBGK)

### 2.8 Statistical analysis

Data are presented as means ± SEM. Proportional variables were transformed using the arcsine square root transformation. Residual normality was confirmed using histograms of raw residuals, plots of residuals vs normal expected value (Q-Q), and plots of residual vs predicted values.

Individual general linear models (GLMs) were fitted for each variable. Treatment and semen group were categorical fixed factors. Type III sums of squares were applied to models with empty cells. To correct for multiple testing, p-values were adjusted using the Benjamini-Hochberg FDR (False Discovery Rate) procedure. Post-hoc of Tukey or Tukey-Kramer (for variables with empty cells) were applied to variables showing significant differences.

Results from YH and WL frozen sperm were analyzed by 2×2 factorial design (YH and WL × fresh and F/T) to evaluate the interaction between breed and treatment for each variable. Due to missing data in some variables, individual factorial analyses were conducted per variable. No significant breed × treatment interactions were found (p > 0.05), so main effects were analyzed independently. P-values from the factorial and the main effects analysis were adjusted using FDR method to account for multiple testing.

Fertility results were analyzed to compare the treatments using Pearson Chi-square tests and FDR adjustments, with a significance set at α = 0.05 for 10 combinations (Fig. 4), 15 combinations (Figure 5A) and 4 combinations (Figure 5B).

**Fig. 4.**
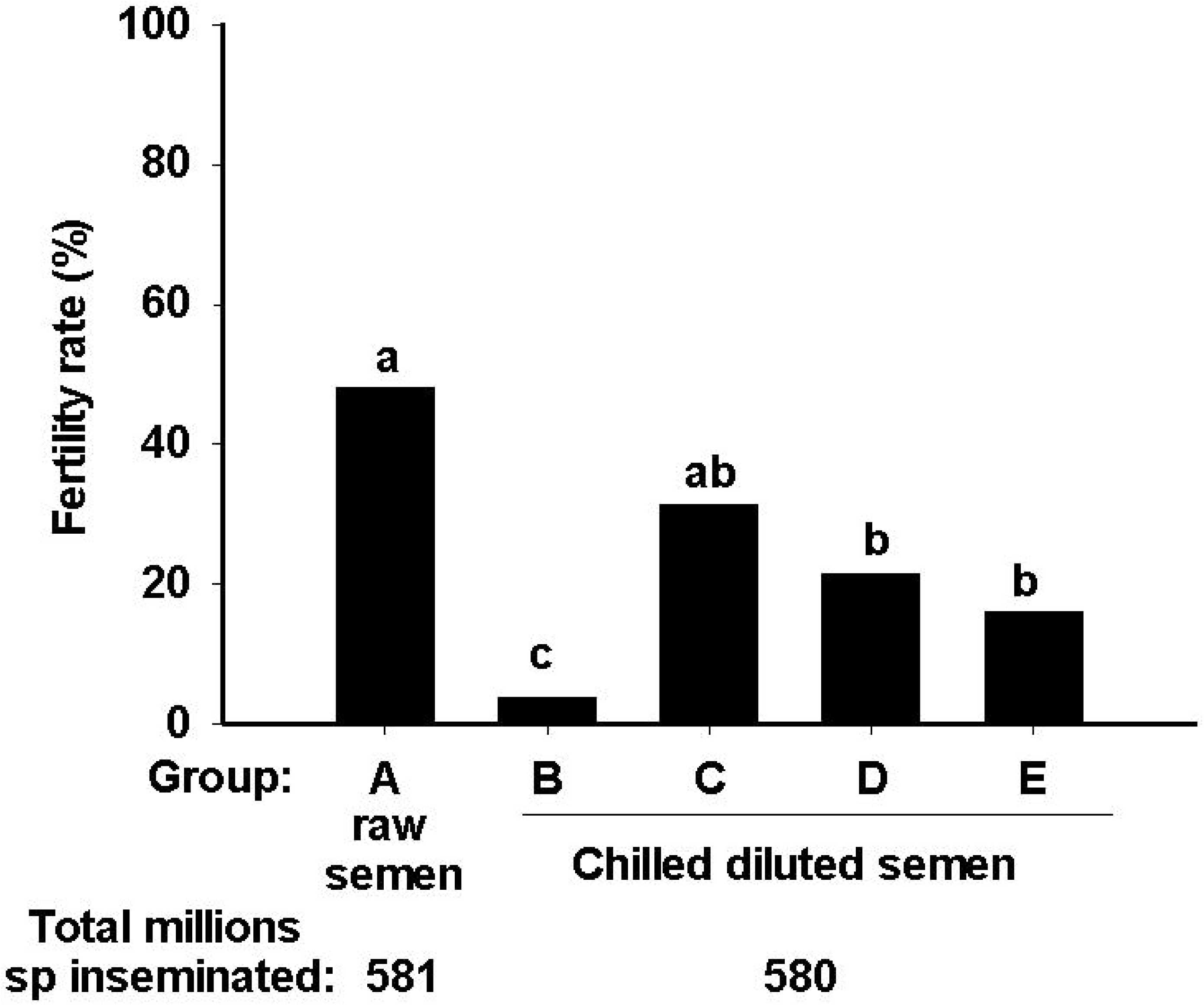
Fertility rates obtained using fresh (not frozen) YH-sp to inseminate WL hens. (A) neat semen; (B) neat semen extended 1:3 in ASG and chilled for 1h at 5°C, C (B+DMA), D (B+AFPI) and E (B+DMA+AFPI). Abbreviations: sp, sperm; YH-sp, Yellow Hungarian sperm, WL, White Leghorn, DMA, dimethylacetamide; AFPI, antifreeze protein I. Different letters on bars represent significant differences (p<0.05) in fertility rate among the groups

**Fig. 5.**
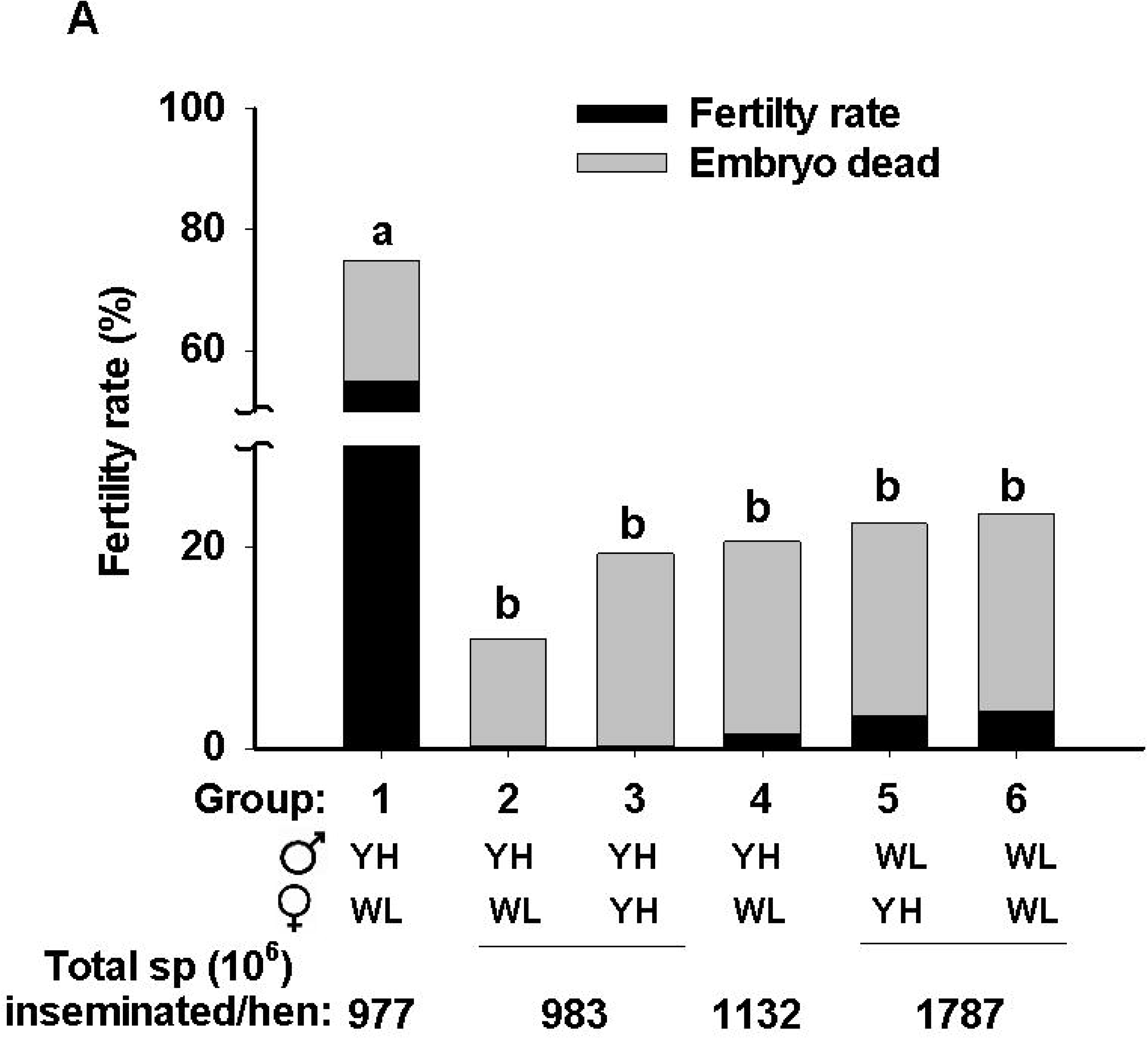
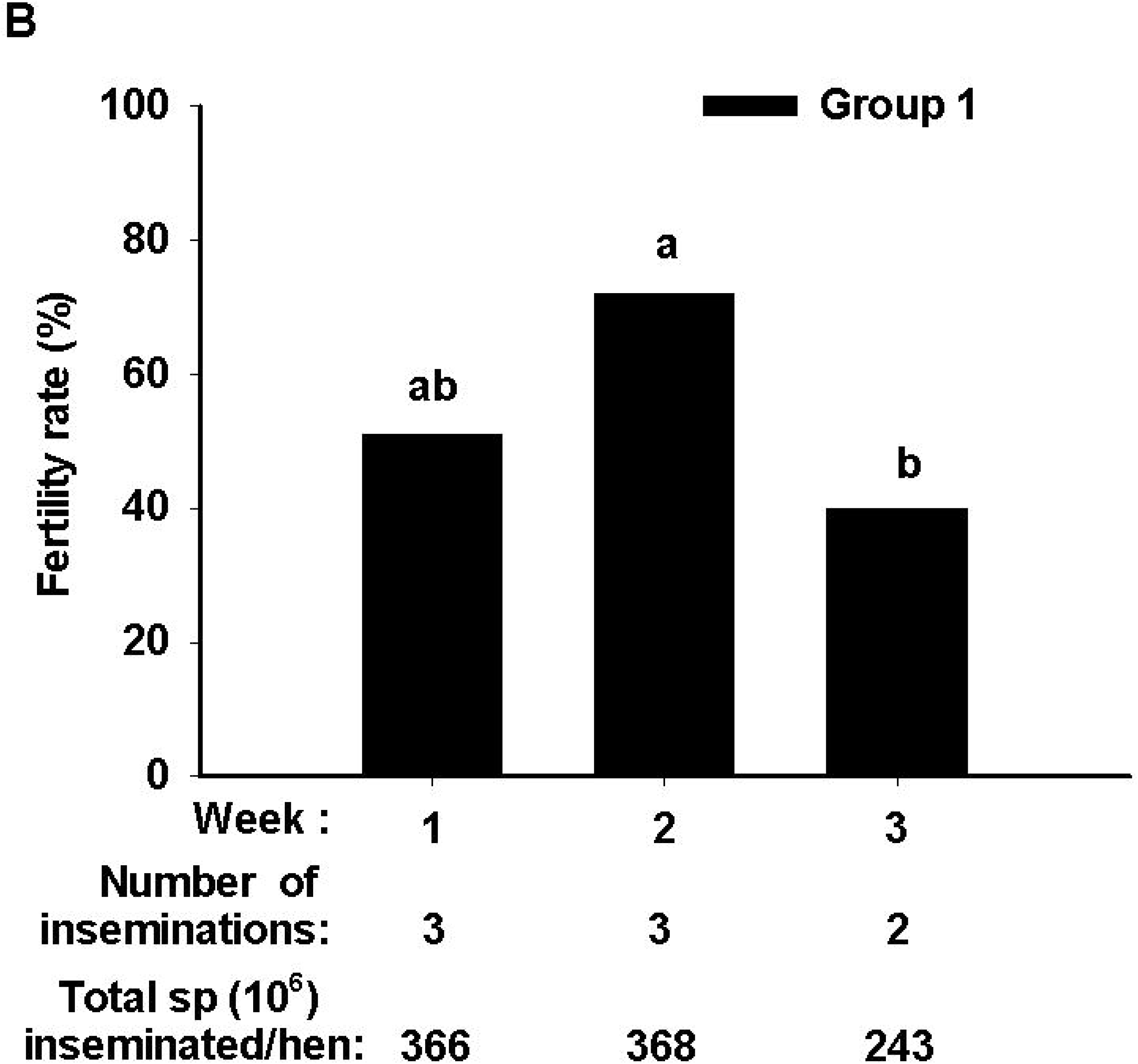
**A.** Black bars: fertility rate obtained from artificial inseminations using raw semen (Group 1) or frozen-thawed sperm (F/T-sp, Groups 2-6) of Yellow Hungarian (YH, Groups 1-4) and White Leghorn (WL, Groups 5-6) roosters (♂) inseminated in YH (Groups 3and 5) and WL (Groups 1, 2,4 and 6) hens (♀). Groups 1-6 corresponds to different combinations of breed and sex: (1) YH-sp raw semen- WL hen; (2) F/T YH- sp - WL hen; (3) F/T YH-sp - YH hen; (4) F/T YH-sp + AFPI - WL hen; (5) F/T WL-sp -YH hen; (6), F/T WL-sp - WL hen. Semen samples used in groups 2-6 were frozen with DMA 0.6 M, but group 4 was additionally supplemented with AFPI (1µg/mL). Grey bars: very early (oviductal) embryonic death identified in eggs initially deemed infertile by candling but later confirmed as fertile through PI staining. **5B.** Fertility rate per week of Group 1. Abbreviations: DMA, dimethylacetamide; AFPI, anti-freeze protein I. Different letters on bars represent significant differences (p<0.05) in fertility rate among the groups or weeks.

All statistical analyses were conducted using TIBCO Statistica™ software, version 13.3 (TIBCO® Software Inc., Palo Alto, CA, USA). Differences were considered statistically significant at P < 0.05.

## 3. RESULTS

Figure 2A illustrates the ability of AF(G)Ps to inhibit ice recrystallization over time, despite being diluted in the poultry extender containing DMA. Smaller ice crystal radii were observed with increasing protein concentrations (Fig. 2A and B). As shown in figures 2A and B, all concentrations of AF(G)Ps tested in the present study exhibited IRI activity.

Freezing of BB sperm revealed no significant differences when supplementing AFGP to the freezing medium. In the case of AFPIII, statistical analysis revealed significant differences (p<0.05) in total motility (TM) and progressive motility (PM) between the control group and all treatments containing AFPIII. However, these differences were not visually evident when comparing the group means (Table 1).

**Table 1.**
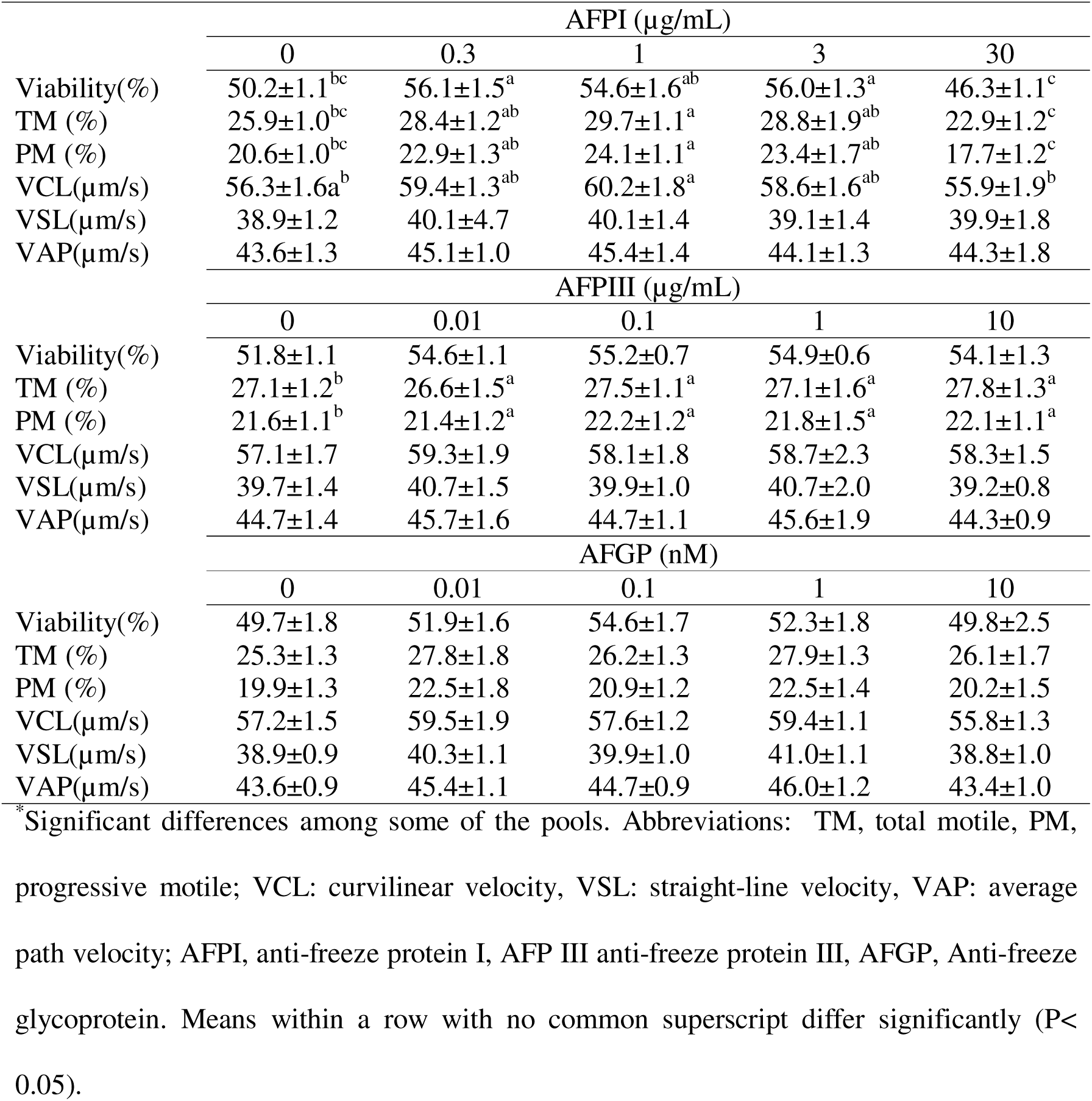
Rooster sperm variables of thawed semen of broiler-breeder roosters, from Netherlands frozen in presence of different anti-freeze (glycol)proteins.

The presence of AFPI increased the viability at concentrations of 0.3 and 3 µg/mL (P=0.007 and 0.009, respectively) (Table 1) while 1 µg/mL increased TM (P=0.03), PM (P=0.034) and VCL (P=0.024).

The results obtained from the three freezing protocols for YH-sp (Table 2) showed significant differences (P<0.05) regarding viability, ILS, and Abn. Protocol 1 gave the highest viability compared to protocol 3 (P=0.0002) and % of ILS compared to Protocol 2 (P=0.0002). Protocol 3 gave the lowest Abn (P=0.0013) compared with Protocol 2. Considering these results, Protocol 1 was selected as the best protocol to test AFPI. The low sperm concentration resulting from the 4x sperm dilution (≈100 million sperm/mL) required an increased number of artificial inseminations from 4 to 8 to adequately assess the fertility of the frozen/thawed (F/T) samples.

**Table 2.** Rooster sperm variables of fresh and frozen-thawed semen of Yellow Hungarian roosters using three different cryopreservation methods.

|  | Fresh | Freezing Method |  |  |
| --- | --- | --- | --- | --- |
|  |  | 1 | 2 | 3 |
| Viability (%)<sup>*</sup> | 92.9±0.9<sup>a</sup> | 56.7±3.0<sup>b</sup> | 49.0±3.4<sup>bc</sup> | 42.8±4.7<sup>c</sup> |
| ILS (%)<sup>*</sup> | 82.3±0.9<sup>a</sup> | 34.5±3.0<sup>b</sup> | 27.7±1.4<sup>c</sup> | 29.7±1.6<sup>bc</sup> |
| Abn (%)<sup>*</sup> | 10.7±0.9<sup>c</sup> | 15.5±1.0<sup>ab</sup> | 19.1±1.6<sup>a</sup> | 12.0±1.4<sup>bc</sup> |
| TM (%)<sup>*</sup> | 65.5±5.4<sup>a</sup> | 27.3±3.4<sup>b</sup> | 26.3±1.8<sup>b</sup> | 21.0±2.7<sup>b</sup> |
| PM (%) | 41.0±7.0<sup>a</sup> | 15.3±3.1<sup>b</sup> | 15.3±1.7<sup>b</sup> | 11.5±2.1<sup>b</sup> |
| VCL(µm/s)<sup>*</sup> | 68.8±6.1<sup>a</sup> | 51.2±2.0<sup>b</sup> | 50.0±4.9<sup>b</sup> | 46.6±2.2<sup>b</sup> |
| VSL(µm/s) | 44.8±5.0<sup>a</sup> | 28.1±2.1<sup>b</sup> | 27.8±2.8<sup>b</sup> | 24.1±2.2<sup>b</sup> |
| VAP(µm/s) | 31.7±3.8<sup>a</sup> | 18.6±2.0<sup>b</sup> | 18.4±1.9<sup>b</sup> | 15.8±2.1<sup>b</sup> |
| AcI(%) | 87.9±1.6<sup>a</sup> | 73.1±4.2<sup>b</sup> | 73.1±4.5<sup>b</sup> | 70.3±5.8<sup>b</sup> |
| DNA fragmentation (%) | ND | 39.2±4.8 | 45.6±3.3 | 49.8±3.3 |

<sup>\*</sup>Significant differences among some of the pools. Abbreviations: ILS, intact live sperm;
Abn, abnormalities, TM, total motile, PM, progressive motile, VCL, curvilinear velocity;
VSL, straight-line velocity; VAP, average path velocity; AcI, acrosome integrity; ND,
not determined. Means within a row with no common superscript differ significantly (P< 0.05).

Sperm samples for AI of the pre-freezing experiment (groups A-E) were characterized before insemination. The results (Table 3) showed higher values for PM, VCL, VSL, VAP in semen of Groups B (P=0.002, 0.002, 0.0003 and 0.0002, respectively) and D (P= 0.02, 0.02, 0.03, 0.01, respectively) compared with Group A. Additionally semen of Group B also had higher TM than Group A (P=0.03). Comparisons were also made between treatments B–E and the original pool of RS from which they were derived (RS for B-E). The results (Table 3) showed a significant decrease in ILS in semen of Group C (semen containing DMA alone) compared to RS for B-E (P=0.02), while the Abn values increased in Groups B (P=0.03), C (P=0.0004) and E (P=0.001). Semen groups C and E (both containing DMA) also showed the lowest values for PM, VSL, and VAP, differing significantly (p<0.05) from Group B, which presented the highest values. Semen of Group C displayed the lowest TM value, significantly lower than those of groups B (P=0.02) and D (P=0.04). No significant differences were observed in viability, VCL, acrosome integrity or DNA integrity among groups.

**Table 3.** Yellow Hungarian sperm variables of: raw semen used for artificial insemination (A, RS for AI)), raw semen aliquoted to prepare treatments B, C, D and E (Raw semen for B-E), cooled extended semen (B), cooled extended semen containing DMA(C), cooled extended semen containing AFPI (D), cooled extended semen containing DMA+AFPI (E). Semen of groups A, B, C, D and E were inseminated (100 million sperm) to White Leghorn hens.

|  | A<br>(RS for AI) | RS for<br>B-E | B<br>(ASG-PE,<br>5°C, 1h) | C<br>(B+DMA) | D<br>(B+AFPI) | E<br>(B+DMA+AF<br>PI) |
| --- | --- | --- | --- | --- | --- | --- |
| Viability (%) | 85.5±3.8 | 93.4±1.1 | 87.5±2.9 | 90.4±1.6 | 86.7±2.1 | 89.1±2.4 |
| Intact live sperm (%) | 81.0±1.2 | 85.7±1.1<sup>a</sup> | 81.4±3.5<sup>ab</sup> | 75.8±2.6<sup>b</sup> | 81.6±2.0<sup>ab</sup> | 77.9±3.7<sup>ab</sup> |
| Abnormalities (%) | 14.1 ±1.7 | 7.8±1.3<sup>c</sup> | 13.6±2.8<sup>ab</sup> | 18.8±2.0<sup>a</sup> | 12.1±2.2<sup>bc</sup> | 18.3±3.4<sup>a</sup> |
| Total motility (%) | 47.7±12.4<sup>BC</sup> | 68.4±10.2<sup>ab</sup> | 80.0±3.4<sup>a, A</sup> | 43.3±6.3<sup>b, C</sup> | 77.4±3.5<sup>a, AB</sup> | 51.2±4.4<sup>ab, ABC</sup> |
| Progressive motility (%) | 19.8±7.3<sup>C</sup> | 42.9±15.8<sup>ab</sup> | 67.9±6.7<sup>a, A</sup> | 27.1±6.2<sup>b, BC</sup> | 55.3±7.5<sup>ab, AB</sup> | 23.9±2.3<sup>b, C</sup> |
| VCL(µm/s) | 55.7±5.9<sup>C</sup> | 81.7±21.6 | 105.7±5.1<sup>A</sup> | 61.7±11.5<sup>C</sup> | 93.5±13.3<sup>AB</sup> | 72.2±13.0<sup>BC</sup> |
| VSL(µm/s) | 20.7±2.4<sup>C</sup> | 37.3±10.6<sup>ab</sup> | 60.7±2.4<sup>a, A</sup> | 24.1±2.3<sup>b, BC</sup> | 40.9±5.2<sup>ab, B</sup> | 27.4±4.9<sup>b, BC</sup> |
| VAP(µm/s) | 29.7±2.6<sup>C</sup> | 51.6±14.9<sup>ab</sup> | 77.4±2.5<sup>a, A</sup> | 35.2±4.0<sup>b, C</sup> | 58.0±6.1<sup>ab, AB</sup> | 40.7±6.7<sup>b, BC</sup> |
| Acrosome integrity (%) | 86.8±5.5 | 89.3±2.6 | 83.5±6.6 | 86.9±2.6 | 84.1±4.1 | 77.6±6.1 |
| DNA integrity (%) | 98.1±1.5 | 96.2±1.7 | 98.7±1.2 | 97.8±0.8 | 97.2±2.6 | 98.4±0.7 |

Different uppercase superscripts indicate significant differences ( $p < 0.05$ ) among “RS for AI” (A) and groups B, C, D and E. Different lowercase superscripts indicate significant differences ( $p < 0.05$ ) among “RS for B-E” and groups B, C, D and E.

AI with treatments A-E revealed significant differences among the groups. Group A showed the highest fertility and had significant differences (P<0.01) with all groups except with Group C (P=0.1). Unexpectedly, a marked decrease in fertility (P=0.0000) was observed (Fig. 4) in Group B (chilled semen in ASG-PE) compared to Group A while Groups C, D, and E exhibited higher fertility rates (P=0.0000, 0.002, 0.002, respectively) than Group B.

Characterization of F/T sperm from YH and WL before AI (Table 4) showed no interaction between freezability and rooster breed. DNA integrity was the only variable affected (P=0.0004)) by rooster breed, with WL semen showing more DNA cryoresistance. The treatment (fresh vs. frozen) resulted in significantly lower values for the frozen/thawed (F/T) samples across all variables, except for VCL (P=0.054). Regarding YH-sp samples frozen with or without AFPI (Table 5), no appreciable effect of AFPI supplementation was seen compared to the non-supplemented samples, except for DNA integrity, which was significantly increased (P=0.004) in the presence of AFPI.

**Table 4.**
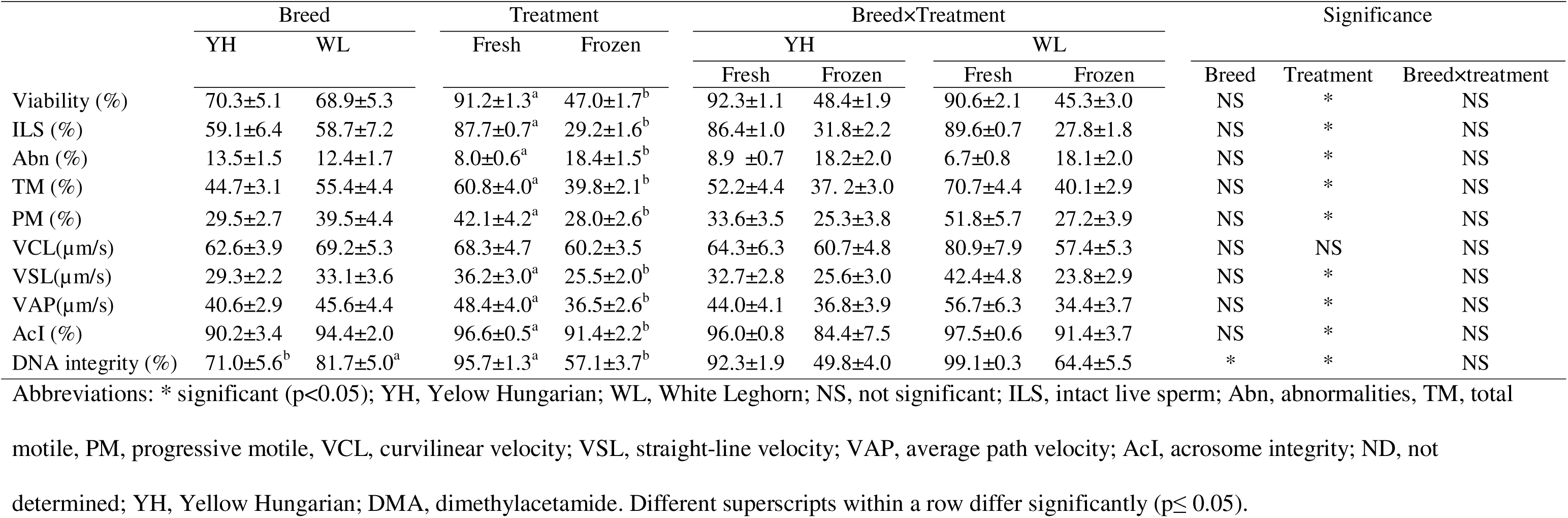
Rooster sperm variables of fresh and frozen-thawed semen of Yellow Hungarian and White Leghorn roosters frozen with DMA.

**Table 5.**
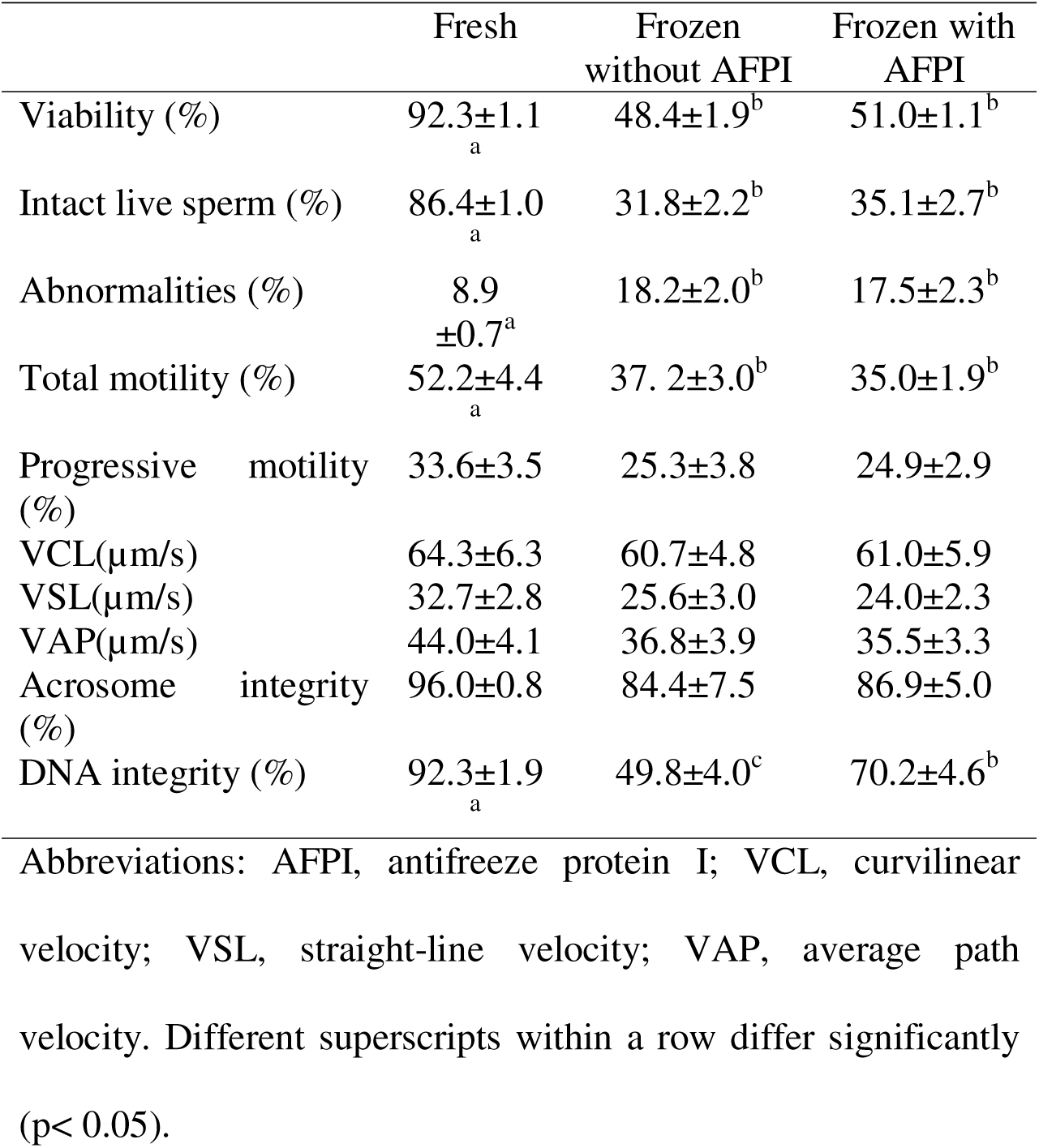
Rooster sperm variables of fresh and frozen-thawed semen of Yellow Hungarian frozen with DMA, in presence or absence of AFPI.

|  | Fresh | Frozen<br>without AFPI | Frozen with<br>AFPI |
| --- | --- | --- | --- |
| Viability (%) | 92.3±1.1<sub>a</sub> | 48.4±1.9<sup>b</sup> | 51.0±1.1<sup>b</sup> |
| Intact live sperm (%) | 86.4±1.0<sub>a</sub> | 31.8±2.2<sup>b</sup> | 35.1±2.7<sup>b</sup> |
| Abnormalities (%) | 8.9<br>±0.7<sup>a</sup> | 18.2±2.0<sup>b</sup> | 17.5±2.3<sup>b</sup> |
| Total motility (%) | 52.2±4.4<sub>a</sub> | 37.2±3.0<sup>b</sup> | 35.0±1.9<sup>b</sup> |
| Progressive motility (%) | 33.6±3.5 | 25.3±3.8 | 24.9±2.9 |
| VCL(µm/s) | 64.3±6.3 | 60.7±4.8 | 61.0±5.9 |
| VSL(µm/s) | 32.7±2.8 | 25.6±3.0 | 24.0±2.3 |
| VAP(µm/s) | 44.0±4.1 | 36.8±3.9 | 35.5±3.3 |
| Acrosome integrity (%) | 96.0±0.8 | 84.4±7.5 | 86.9±5.0 |
| DNA integrity (%) | 92.3±1.9<sub>a</sub> | 49.8±4.0<sup>c</sup> | 70.2±4.6<sup>b</sup> |

Abbreviations: AFPI, antifreeze protein I; VCL, curvilinear velocity; VSL, straight-line velocity; VAP, average path velocity. Different superscripts within a row differ significantly (p< 0.05).

The results obtained from artificial insemination (AI) with F/T sperm are shown in Figure 5A. No fertility was observed in Groups 2 and 3. Group 4, which received AFPI, achieved a fertility rate of 1.5%, though this was not significantly different from the other groups. Groups 5 and 6, inseminated with frozen/thawed (F/T) sperm from White Leghorn (WL) roosters, showed fertility rates of 3.4% and 3.9%, respectively. No significant differences were observed with respect to the breed of hen.

Closer inspection of the eggs initially classified as unfertile during candling revealed that approximately 18% had been fertilized (Fig. 5A), but the embryos had died at very early stage, likely within the oviduct.

Analysis of the fertility rate per week (Fig. 5B) of raw semen (Group 1) showed that the maximum fertility rate could not be maintained in the 3^rd^ week, when the number of inseminations was decreased to 2 per week.

## 4. DISCUSION

Comparisons among the different AF(G)Ps revealed that AFPI exhibited a superior protective effect compared to AFPIII and AFGP, despite all tested concentrations of these proteins exhibiting IRI activity. In this respect, research has demonstrated that performance of individual AF(G)Ps may vary depending on factors such as their adsorption rate, cooling rate or the degree of supercooling experienced [13]. Moreover, studies using model membranes with varying lipid types demonstrated that AFPI and AFGP interactions are lipid-specific, indicating that the protective efficacy of these proteins is influenced by the composition of the lipid bilayer [34,35].

For the YH semen, we evaluated both the DR 4× used with BB-sp and DR 2.3×. These treatments were further compared to a standard freezing protocol for YH semen, which employs a DR of 2.1×. The results showed that the DR 4× yielded the highest viability. The beneficial effect of higher DRs compared to lower ones have been also reported in other studies [2,36]. A possible explanation could be the dilution of harmful substances present in raw semen [5,37,38], and/or the presence of beneficial components in the diluent—excluding DMA, as all three DRs used the same final DMA concentration 0.6 mol/L.

Sperm characterization of the 5 pre-freezing steps (groups A-E) showed that chilled sperm with DMA had lower values in most of the motility parameters (except TM) compared to ASG-PE alone, which yielded the highest motility—even higher than the “RS for AI” group. Based on these results, better fertility was expected with semen chilled in ASG-PE and poorer outcomes with the semen chilled in presence of DMA. However, fertility decreased in all groups compared to the “RS for AI” group. Surprisingly, adding DMA to ASG-PE strongly mitigated this decline, eliminating the significant differences in fertility compared with the “RS for AI” group.

AFPI alone prevented significantly the increase in sperm abnormalities, in contrast to DMA and DMA-AFPI treatments, which showed the highest abnormality rates. Despite this significant difference, AFPI and DMA-AFPI exhibited similar fertility rates. AFPI and AFPI-DMA treatments also showed reduced fertility versus the “RS for AI” group, but their fertility rates remained significantly superior to that of ASG-PE alone

Among the possible causes that explain the loss of fertility observed with chilled sperm in ASG-PE we could find, a detrimental effect of the ASG-PE itself, or damage induced by cooling/chilling that was not detected by the evaluated parameters. The first hypothesis was discarded based on findings from a separate study (not part of this work), in which insemination with ASG-PE treated sperm, but not subjected to chilling, resulted in higher fertility rates than chilled semen [39] which points chilling injury as a possible cause of fertility loss.

No serious changes in sperm motility, viability or acrosome integrity were found across treatments A-E, however the serious impact of chilling on the fertility was evident after AI. This could be related to damages induced by rewarming [40] of the chilled sperm inside the hen or to damages that are only evident after rewarming.

A similar loss of chicken sperm fertility after chilling in absence of cryoprotectant, was previously reported by Elomda et al., (2024) [41]; however, the impact on fertility was less pronounced (18.2%), which coincided, in their study, with the presence of polyvinylpyrrolidone (PVP) in the chicken extender. PVP has been hypothesized to exert cryoprotective effects through colligative properties, by coating cell membranes [42] or by inhibiting freezing-induced protein dissociation [43]. Thus, supplementing protective macromolecules in the poultry extender may perhaps prevent the loss of fertility during cooling/chilling.

The loss of fertility observed during chilling was significant mitigated by the presence of DMA or AFPI, observing the highest protection role for the former. The protective role of internal cryoprotectants as DMA is attributed to their capacity to maintain the fluidity of the sperm membranes at low temperatures [44] by decreasing the lipid phase transition temperature of the cell membranes. Regarding AFPI, several works have reported a stabilizing effect of AF(G)Ps of the cell membranes [34,45]. There was no additional gain by using the AFPI-DMA combination.

Artificial insemination with F/T sperm of YH gave zero fertility in absence of the AFPI, independently of the breed of hen, and some very low fertility in the presence of the protein. Inseminations with RS ruled out infertility in the YH roosters. However, it is worth noting that, after four inseminations with RS in the current batch of hens, the fertility rate was only about 50%, suggesting limited reproductive performance within this group.

The fertility rates obtained with F/T sperm could suggest that the freezing protocol failed to preserve the functionality of the sperm. However, the analysis on embryo dead in the fertility assays with F/T sperm indicated that a percentage of sperm, from YH and WL roosters, were able to fertilize the eggs but the embryos were not able to continue developing. This observation could be a consequence of inseminating an insufficient number of spermatozoa, caused by the low concentration of semen doses. According to Hemmings and Birkhead (2015) [46], when very few sperm penetrate the avian ovum, embryos are unlikely to survive beyond the earliest stages of development. The inability to maintain fertility over time with raw semen doses may reflect limited sperm storage in the sperm storage tubules (SSTs) compromising the embryo development. On this basis, the observation that hens (YH and WL) inseminated with WL sperm showed higher fertility rates (though not statistically significant) compared to those inseminated with YH sperm could be attributed to the higher sperm dose of the WL group which contained approximately 1.8 times more spermatozoa. However, the higher DNA cryoresistance observed in WL sperm may have also contributed to these results.

In conclusion, cryopreservation of semen could be an instrument in conserving rare breeds, but freezing and thawing of YH semen is still a challenge.

This study showed that pre-freeze processing steps such as extending-cooling, can already strongly reduce the fertility of the semen, albeit, this reduction was lower in the presence of DMA and/or AFPI. Future studies may address why the pre-freeze fertility was lowered and how it can be prevented, e.g. by using protective macromolecules already during cooling, and/or by more gradual cooling.

With the current freezing protocol and insemination dose, the fertility of frozen/thawed (F/T) semen was very low. Given that sperm concentration in raw YH semen is also relatively low, it becomes necessary to increase the sperm dose without compromising post-thaw sperm quality.

AFPI had a small but significant positive effect on some post-thaw sperm variables. It is yet unclear whether AFPI may also affect fertility of F/T semen, albeit that the only fertile eggs obtained with F/T YH semen were obtained using AFPI. It may be necessary to change pre-freeze semen processing to avoid pre-freeze loss of fertility and to yield sufficiently concentrated insemination sperm dosages.

## ACKNOWLEDGEMENTS

This work was funded by the European Union, project 101109983 — CRYOCHICK — HORIZON-MSCA-2022-PF-01, Ministry of LVVN, project KB-34-013-002, the Dutch Research Council to T.P.H (NWO-FoI.2024.016) and I.K.V. (NWO-VICI 232.106) and the European Research Council to I.K.V. (ERC-2020-CoG 101001965).

## DECLARATION OF GENERATIVE AI AND AI-ASSISTED TECHNOLOGIES IN THE WRITING PROCESS

During the preparation of this work the author(s) used ChatGPT in order to improve language and readability. After using this tool, the author(s) reviewed and edited the content as needed and take(s) full responsibility for the content of the publication.

